# Chromosome-scale assembly and annotation of the wild wheat relative *Aegilops comosa*

**DOI:** 10.1101/2024.10.15.618371

**Authors:** Hongna Li, Shams ur Rehman, Rui Song, Liang Qiao, Xiaohua Hao, Jing Zhang, Kairong Li, Lifeng Hou, Wanyi Hu, Le Wang, Shisheng Chen

**Affiliations:** National Key Laboratory of Wheat Improvement, Peking University Institute of Advanced Agricultural Sciences, Shandong Laboratory of Advanced Agriculture Sciences in Weifang, Weifang 261325, Shandong, China; State Key Laboratory of Cellular Stress Biology, School of Life Sciences, Xiamen University, Xiamen 361102, China; College of Plant Science, Jilin University, Changchun 130062, China

## Abstract

Wild relatives of wheat are valuable sources for enhancing the genetic diversity of common wheat. *Aegilops comosa*, an annual diploid species with an MM genome constitution, possesses numerous agronomically valuable traits that can be exploited for wheat improvement. In this study, we report a chromosome-level genome assembly of *Ae. comosa* accession PI 551049, generated using PacBio high-fidelity (HiFi) reads and high-throughput chromosome conformation capture (Hi-C) data. The assembly spans 4.47 Gb, featuring a contig N50 of 23.59 Mb and a scaffold N50 of 619.05 Mb. A total of 39,063 gene models were annotated through a combination of homoeologous proteins, Iso-Seq, and RNA-Seq data. Comparative genome analysis revealed a terminal intrachromosomal translocation in chromosome 2M of *Ae. comosa* (and *Ae. umbellulata*) compared to its homoeologous chromosomes in other diploid wheat species. Phylogenetic analysis showed a close relationship between *Ae. comosa* and *Ae. umbellulata*. This newly constructed reference genome of *Ae. comosa* will serve as an important genomic resource for comparative genomic studies and the cloning of agriculturally important genes.

## Background & Summary

*Aegilops comosa* (2n = 2x = 14, MM), is an annual diploid species belonging to the tertiary gene pool of wheat. It predominantly occurs in the coastal regions of Albania, the former Yugoslavia, and both coastal and inland areas of Greece^1-3^. As a diploid M genome donor species, *Ae. comosa* plays a crucial role in the evolution of polyploid *Aegilops* species, including *Ae. biuncialis* (U^b^U^b^M^b^M^b^) and *Ae. geniculate* (U^g^U^g^M^g^M^g^)^1,4^. Both diploid and polyploid M genome species exhibit robust resistance to various diseases such as stem rust^5^, stripe rust^3,6^, leaf rust^7^, and powdery mildew^6^, as well as resistance to multiple pests^3^. Additionally, the M genome harbors novel low-molecular-weight glutenin subunits and genes associated with salt tolerance^8^. However, the discovery and utilization of beneficial genes from *Ae. comosa* have been greatly limited thus far due to the absence of a reference genome and comprehensive annotation for this species.

The *Ae. comosa* accession PI 551049 has previously been documented to exhibit strong resistance against all tested races of *Puccinia graminis* f. sp. *tritici* (*Pgt*) including TTKSK (Ug99), TRTTF, TTTTF, TPMKC, RKRQC, QTHJC, and QFCSC^9^.

In this study, PI 551049 also displayed high resistance to the Chinese *Pgt* race 34MKGQM (Fig. 1a). To facilitate future research, we assembled a chromosome-scale reference genome for *Ae. comosa* accession PI 551049, using a combination of sequencing technologies: 276.84 Gb HiFi reads (∼62 ×), 473.44 Gb Hi-C reads (∼105 ×), and 228.69 Gb whole-genome sequencing (WGS) short reads (∼51 ×) (Table 1). The initial genome assembly, generated using the *hifiasm*^10^ assembler with HiFi reads. After filtering out mitochondrial, chloroplast, and microbial sequences, the primary contigs were anchored into pseudochromosomes using *3D-DNA* and *Juicer* with Hi-C data. The final genome assembly comprised 4.47 Gb, with a contig N50 of 23.59 Mb and a scaffold N50 of 619.05 Mb. Impressively, 97.49% of the assembly was assigned to seven pseudo-chromosomes (Fig. 1b). The assembly contained 336 gaps, and ten telomeres were identified using the telomere repeat sequence (“CCCTAAA”) across these seven chromosomes (Table 2). The quality of the assembly was further validated by a Long Terminal Repeat Assembly Index (LAI) score of 17.67, a Benchmarking Universal Single-Copy Orthologs (BUSCO) score of 97.30% using the poales_odb10 database, and a consensus quality value (QV) score of 43.23, all reflecting the high accuracy and completeness of the genome assembly.

**Table 1.**
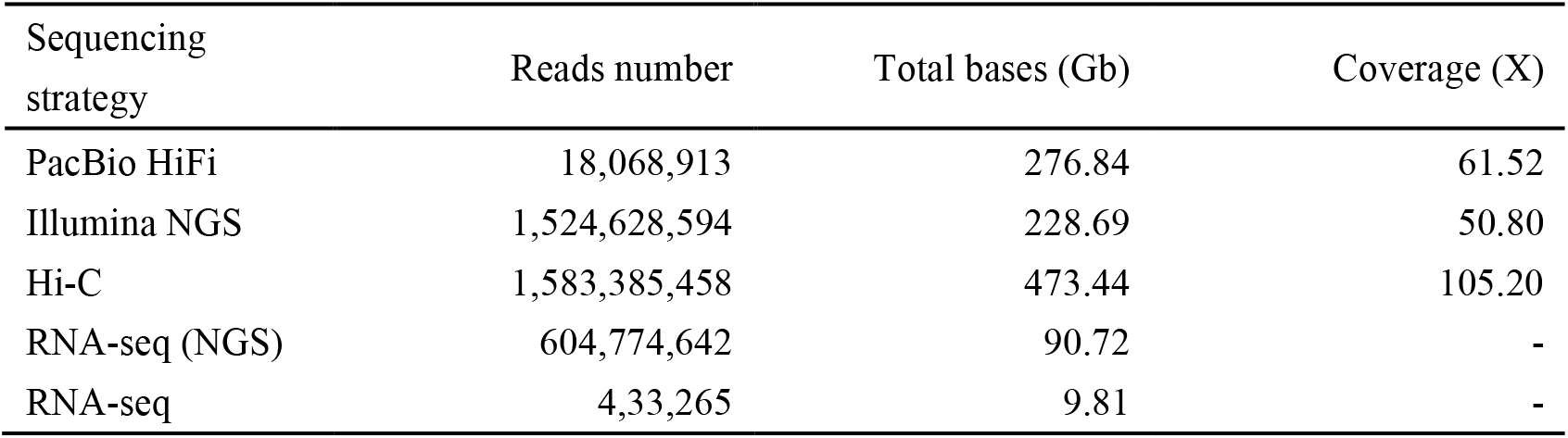
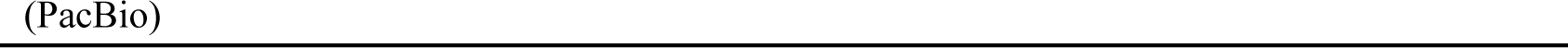
Sequencing data used for *de novo* genome assembly and annotation.

**Table 2.** Summary of chromosome information of the *Ae. comosa* genome.

| Chromosome | Length (bp) | Percentage (%) | Gap number | Telomeres number | Gene number | Centromere region |
| --- | --- | --- | --- | --- | --- | --- |
| Chr1M | 557,637,258 | 12.46 | 50 | 1 | 4,338 | ~206-218 Mb |
| Chr2M | 644,874,278 | 14.41 | 52 | 1 | 5,637 | ~183-196 Mb |
| Chr3M | 693,849,849 | 15.50 | 47 | 1 | 5,259 | ~278-289 Mb |
| Chr4M | 574,023,925 | 12.82 | 57 | 1 | 3,668 | ~222-240 Mb |
| Chr5M | 619,056,011 | 13.83 | 40 | 2 | 5,345 | ~194-206 Mb |
| Chr6M | 574,811,486 | 12.84 | 45 | 2 | 4,297 | ~283-296 Mb |
| Chr7M | 699,149,154 | 15.62 | 45 | 2 | 5,262 | ~351-363 Mb |
| Total | 4,363,401,961 | 97.49 | 336 | 10 | 33,806 | - |
| Unplaced | 112,485,487 | 2.51 | - | - | 5,257 | - |

**Fig. 1.**
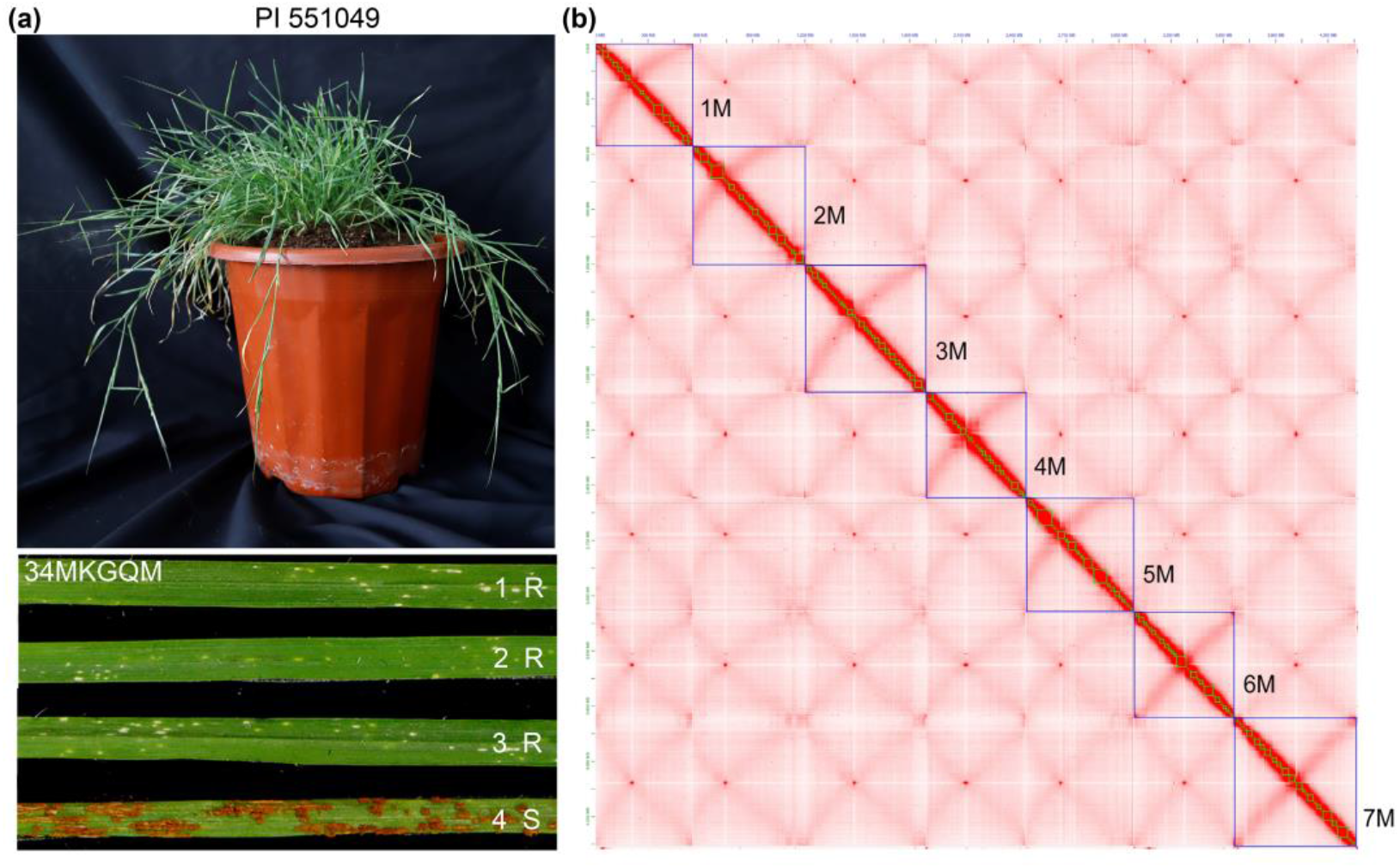
Morphological characteristics and Hi-C interaction heatmap. **(a)** Morphological characteristics of *Ae. comosa* accession PI 551049 and its response to the Chinese *Pgt* race 34MKGQM. *Ae. comosa* accession PI 551054 served as a susceptible control. 1-3, PI 551049; 4, PI 551054. R, resistant; S, susceptible. **(b)** Hi-C interaction heatmap post-integration of Hi-C data and manual correction. Blue boxes represent pseudochromosomes, and green boxes indicate contigs

Genome annotation revealed that the *Ae. comosa* genome contains approximately 3.86 Gb (86.30%) of repetitive sequences, predominantly composed of *Gypsy* (43.73%) and *Copia* (23.47%) LTR retrotransposons (Fig. 2 and Table 3). Given that the *Gypsy* families *RLG_Cereba* and *RLG_Quinta* are enriched in centromeres, we analyzed their distribution within *Ae. comosa* to pinpoint the locations of chromosome centromeres (Fig. 3a). A total of 39,063 protein-coding genes were predicted in *Ae. comosa*, based on homoeologous proteins, RNA-seq and Iso-seq data from four different tissues (leaf, root, stem, and spike) at various developmental stages of PI 551049. The average gene density was around 7.77 genes per Mb, with an average gene length of 3326.32 bp. Of the predicted genes, 37,821 were successfully functionally annotated using the eggNOG-mapper tool (Table 3).

**Table 3.** Statistics of the *Ae. comosa* genome assembly and annotation.

| Genome assembly | Value |
| --- | --- |
| Total assembly size (Gb) | 4.47 |
| GC content (%) | 46.63 |
| N50 of contig (Mb) | 23.59 |
| Number of scaffolds | 722 |
| N50 of scaffold (Mb) | 619.05 |
| Max scaffold size (Mb) | 698.53 |
| Assign rate | 97.49 |
| BUSCO completeness (%) | 97.30 |
| LAI (LTR Assembly Index) | 17.67 |
| QV (Quality Value) | 43.23 |
| **Repetitive sequences (% of genome)** |  |
| SINEs (bp) | 2,258,176 (0.05%) |
| LINEs (bp) | 71,723,038 (1.60%) |
| LTR (bp) | 3,016,172,536 (67.39%) |
| Copia (bp) | 1,050,697,085 (23.47%) |
| Gypsy (bp) | 1,957,218,909 (43.73%) |
| DNA transposons (bp) | 467,286,877 (10.44%) |
| Other | 307,623,120 (6.87%) |
| Total (bp) | 3,862,805,571 (86.30%) |
| **Gene annotation** |  |
| No. of the protein-coding genes | 39,063 |
| No. of genes with egg-NOG annotation | 37,821 (96.82%) |
| No. of genes per Mb | 7.77 |
| The average length of genes | 3326.32 |
| BUSCO completeness (%) | 97.60% |

**Fig. 2.**
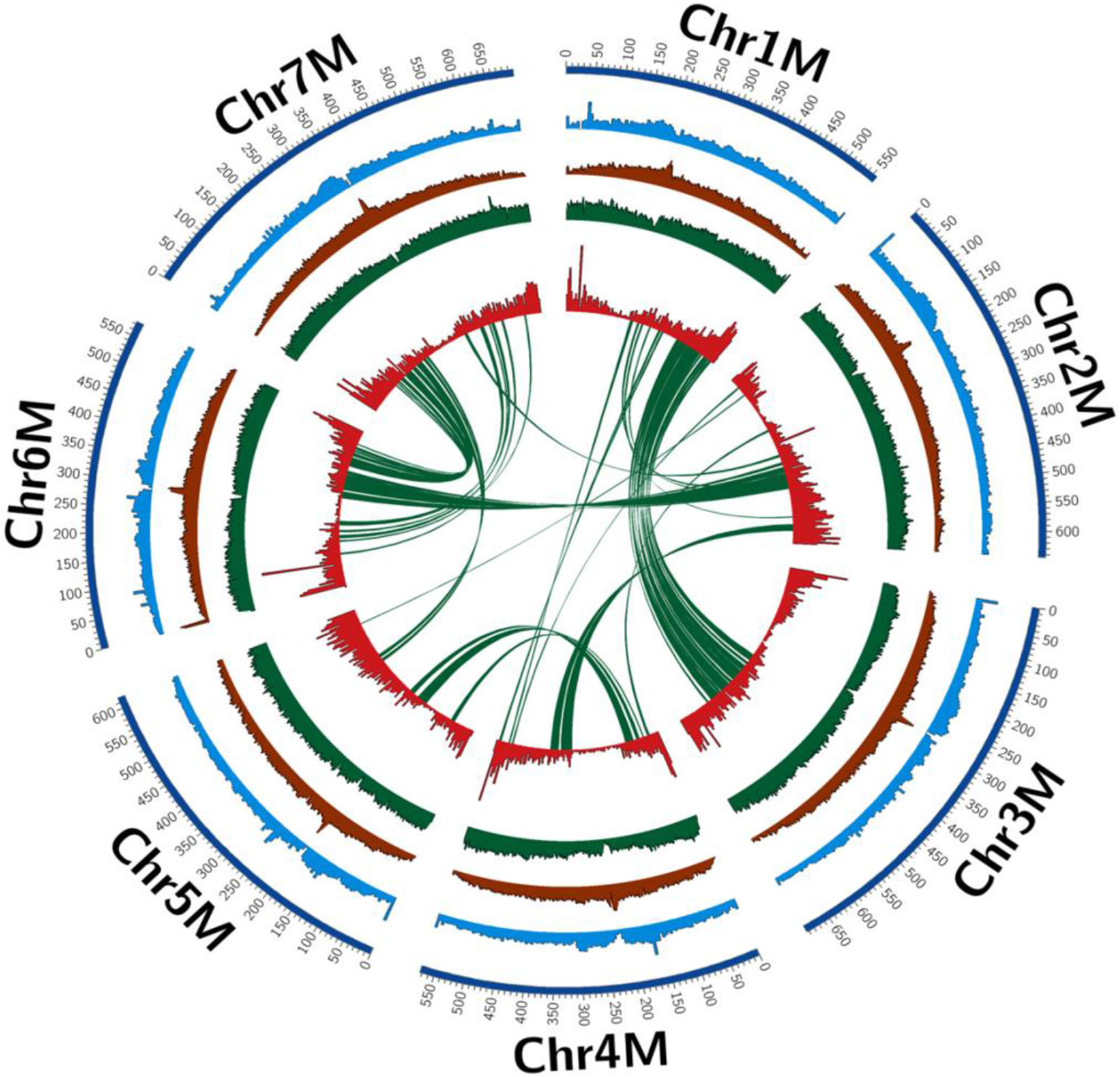
Genomic features of the *Ae. comosa* genome. The tracks, arranged from outermost to innermost, include: pseudochromosomes, GC content, LTR/*Gypsy* density, LTR/*Copia* density, protein-coding genes density, and paralog synteny relationships.

**Fig. 3.**
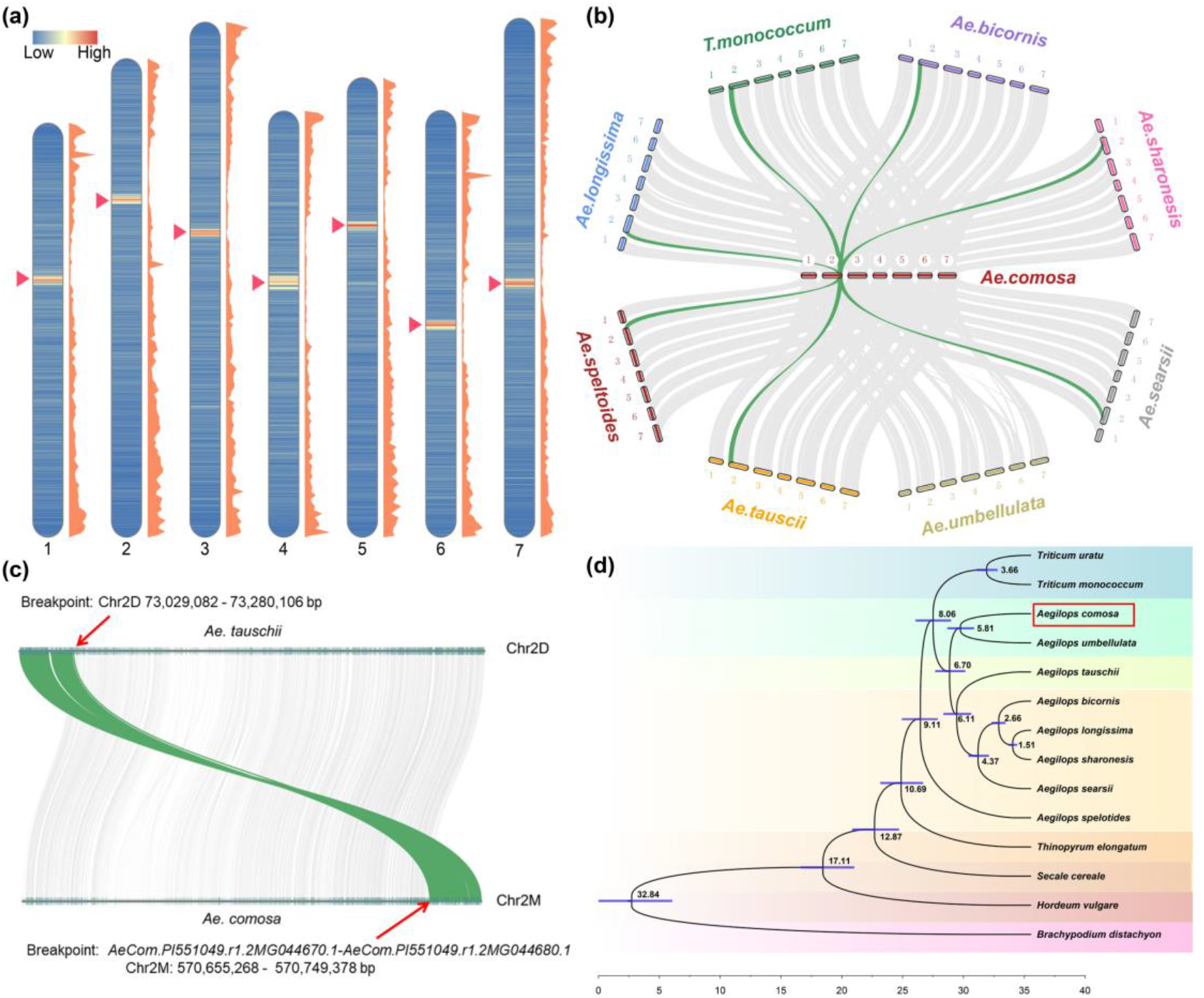
Characterization of the genome of *Ae. comosa* accession PI 551049. **(a)** The density of the *RLG_Cereba* and *RLG_Quinta* was calculated in 1 Mb sliding windows (left) and the density of protein-coding genes was calculated in 5Mb windows (right). Red arrowheads indicate the centromere regions. **(b)** Syntenic analysis between *Ae. comosa* and other diploid species. The green lines indicate the intrachromosomal translocation region in chromosome 2M of *Ae. comosa* compared to its homoeologous chromosomes in other diploid species. **(c)** Syntenic relationships between *Ae. comosa* and *Ae. tauschii* on chromosome 2. Arrows represent the translocation breakpoints. **(d)** Phylogenetic tree of *Ae. comosa* and 13 diploid species based on 2,342 single-copy orthologous genes. The red box indicates the *Ae. comosa* genome.

Synteny analysis demonstrated strong collinearity between the genome of *Ae. comosa* and those of *Triticutum monococcum* (A^m^A^m^), *Ae. tauschii* (DD), *Ae. umbellulata* (UU), *Ae. searsii* (SS), *Ae. sharonesis* (SS), *Ae. bicornis* (SS), *Ae. longissima* (SS), and *Ae. speltoides* (SS) (Fig. 3b). However, detailed analysis of chromosome 2 revealed that the 2M chromosome in *Ae. comosa* has undergone structural rearrangements through a terminal intrachromosomal translocation, distinguishing it from its homoeologous chromosomes in other diploid species, with the exception of *Ae. umbellulata* (Fig. 3b). This finding aligns with the study by Said et al. (2021), who observed a distal segment of the short arm located at the distal end of group 2 long arms in both *Ae. comosa* and *Ae. umbellulata*^11^. Microcollinearity analysis between *Ae. comosa* and *Ae. tauschii* pinpointed the translocation breakpoint between the genes *AeCom.PI551049.r1.2MG044670.1* and *AeCom.PI551049.r1.2MG044680.1* (Chr2M: 570,655,268 - 570,749,378 bp), affecting approximately 74.2 Mb of sequences from 2MS to 2ML (Fig. 3c). A phylogenetic tree, constructed using 2,342 single-copy orthologous genes from *Ae. comosa* and 13 other monocot/*Aegilops* species, with *Brachypodium distachyon* as the outgroup, revealed that *Ae. comosa* is closely related to *Ae. umbellulata* (Fig. 3d). Molecular dating analyses estimate the divergence between *Ae. comosa* and *Ae. umbellulata* occurred around 5.81 million years ago (Fig. 3d).

In summary, this research provides the first chromosome-level reference genome for the M genome of the Triticeae tribe. This genome will serve as a foundation for future studies on M genome introgression into wheat and for the isolation of agriculturally significant genes originating from *Ae. comosa*.

## Methods

### Plant material, DNA extraction, and sequencing

*Ae. comosa* accession PI 551049 was grown in a greenhouse at the Peking University Institute of Advanced Agricultural Sciences, Weifang, China. Fresh leaves of two-week-old plants were selected for whole-genome sequencing. Tissues from the root, stem, leaf, and spike of PI 551049 at various growth stages were collected for RNA-sequencing. Immediately after harvesting, each sample was flash-frozen in liquid nitrogen and stored at -80°C.

High molecular weight (HMW) DNA was extracted from young leaves of PI 551049 using the CTAB protocol^12^. The HMW DNA was used for single-molecule sequencing, whole-genome sequencing, and Hi-C (high-throughput chromosome conformation capture) sequencing. For genome assembly, 15-20 kb libraries were constructed using the SMRTbell Template Preparation Kit 2.0 according to the manufacturer’s instructions, and PacBio HiFi long reads were generated using the PacBio Sequel Revio sequencing platform at Berry Genomics, Beijing, China. For genome surveys, approximately 228.69 Gb (50.80 × coverage) of 150 bp paired-end reads were sequenced on an Illumina NovaSeq 6000. For Hi-C sequencing, libraries were prepared from leaf tissue of PI 551049 by cross-linked using a 2% formaldehyde solution to capture interactions, following a published protocol^13^. After enzymatic digestion with 400 units of *Hind*III, the cross-linked DNA fragments were blunt-end ligated after biotin-14-dCTP was added to the DNA ends. Biotin was removed from non-ligated fragment ends using T4 DNA polymerase. After re-ligation, reverse cross-linking, and purification, the chromatin DNA was sheared to a size of 300-700 bp by sonication. The biotin-labeled fragments were then enriched using magnetic beads. After adding A-tails and sequencing adapters to the fragment ends, 150-bp paired-end reads were generated on the Illumina NovaSeq 6000 during Hi-C library preparation and subsequent analysis.

### Genome survey

Fastp^14^ was used to filter out adaptor sequences, low-quality reads, and short reads from the next-generation sequencing data with the default settings. Next, the frequency distribution of the depth of clean data with 21-mers was computed using Jellyfish v2.2.10^15^. The estimated genome size, heterozygosity ratio, and repeat content distributions were calculated using *find*GSE v0.1.0^16^, determining the genome size to be approximately 4.51 Gb (Fig. 4).

**Fig. 4.**
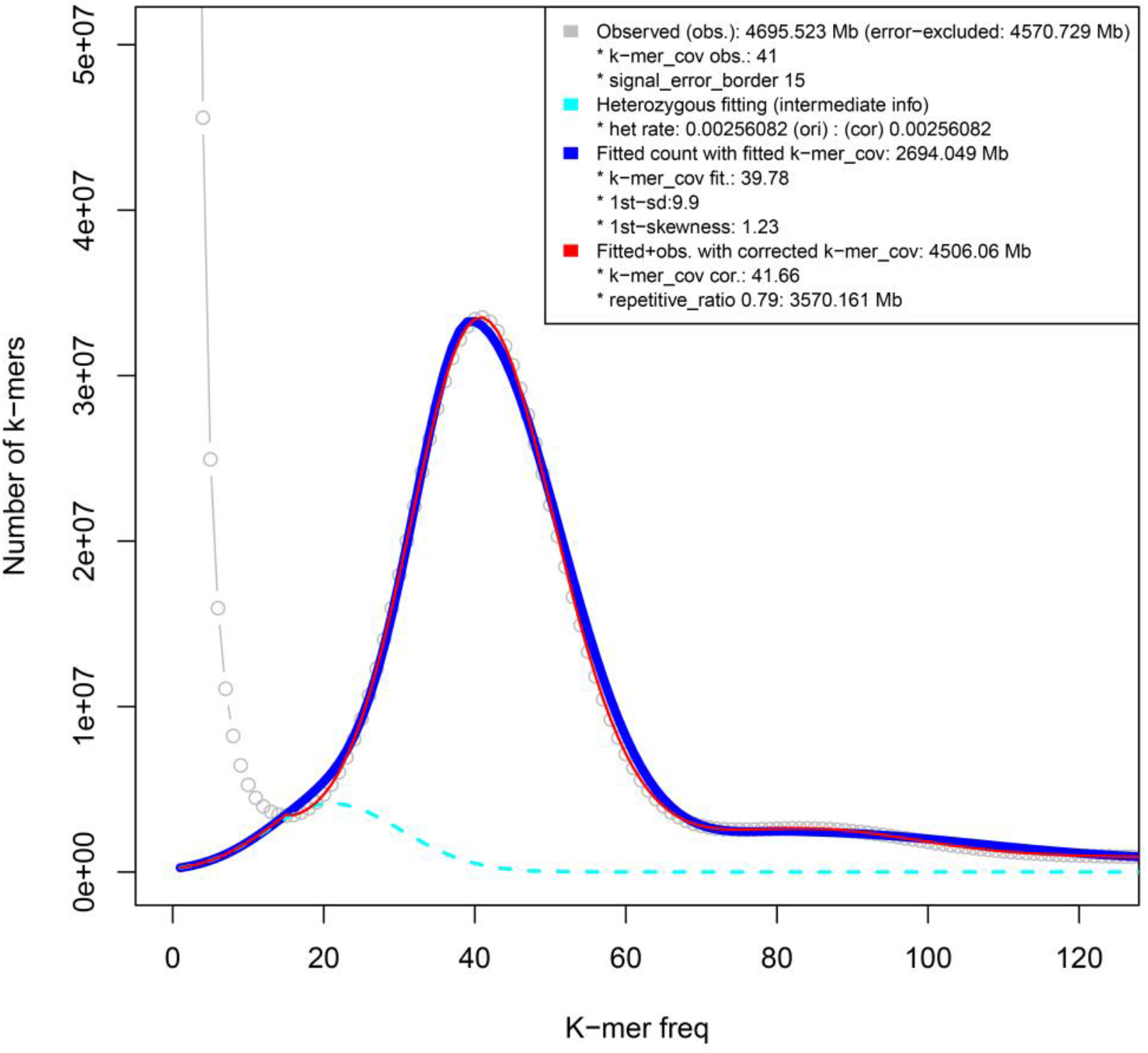
K-mer based estimation of genome characteristics of *Ae. comosa*.

### Chromosome-level assembly

For genome assembly, HiFi long reads were used for *de novo* assembly into contigs using *hifiasm* v0.19.5-r587^10^ with the parameter “-l 0”. Contigs with more than 50% coverage of mitochondrial or chloroplast sequences were identified by aligning them to references genome using minimap2 v2.24-r1122^17^ with parameter “-x asm5” and subsequently removed. To anchor the contigs to chromosomes, Hi-C sequencing data were mapped onto the assembled contigs using the Juicer v1.5^18^ pipeline, with *Hind*III as the enzyme site. Chromosome-level scaffolds were then constructed using the 3D-DNA v190716^19^ pipeline (run-asm-pipeline.sh with the “-r 0” parameter). Finally, scaffold boundaries were manually corrected using JuiceBox v1.11^20^.

### Identification and characterization of repetitive elements

For genome annotation, *de novo* transposable elements (TE) were identified using RepeatModeler v2.0.3^21^ to construct a TE library. Long Terminal Repeat (LTR) retrotransposons were specifically identified with LTR_FINDER_parallel v1.2^22^. All repetitive elements in the genome of *Ae. comosa* were masked using RepeatMasker v4.1.5 (https://www.repeatmasker.org/) with default parameters. A total of 3.86 Gb (86.30%) of the assembled genome was identified as repetitive sequences. Previous studies highlighted the enrichment of the *Gypsy* families *Cereba* and *Quinta* in centromeric regions^23^. To confirm this, LTR/*Gypsy* retrotransposon sequences were aligned with the consensus sequences (RLG_Taes_Cereba_consensus-1 and RLG_Taes_Quinta_consensus-1), obtained from the Transposable Elements Platform (https://trep-db.uzh.ch/) using BLASTN. Finally, the positions of the chromosome centromeres in *Ae. comosa* were identified based on the distribution of *RLG_Cereba* and *RLG_Quinta*.

### Gene prediction and functional annotation

To annotate protein-coding genes in *Ae. comosa*, we used multiple approaches. First, repetitive sequences within the genome were masked to focus on regions likely to contain genes. The repeat-masked genome was then used for annotating gene models. Second, we used a combination of homoeologous proteins, Iso-Seq, and RNA-Seq analysis for protein-coding gene prediction. RNA-seq data obtained from four tissues (leaf, root, stem, and spike) at various developmental stages were mapped to PI 551049 assembly using Hisat2 v2.2.1^24^, and transcripts were assembled with StringTie v2.2.1^24^ (parameters: -f 0.3 -m 150 -t). Iso-Seq raw data were processed using CCS v6.4.0 (https://ccs.how/), lima v2.6.0 (https://lima.how/), and isoseq2 v3.8.2 (https://github.com/PacificBiosciences/IsoSeq), and the resulting transcripts were mapped with minimap2 v2.26-r1175^17^. Redundant isoforms were collapsed into transcript using cDNA_Cupcake (parameters: --dun-merge-5-shorter). The RNA-seq and Iso-Seq transcripts were merged using Stringtie v2.2.1^24^ (parameters: --merge -m 150 -t). Open reading frames (ORFs) were identified within the merged transcripts using Transdecoder v5.5.0 (https://github.com/TransDecoder/TransDecoder). For homoeologous protein data, annotations from *Ae. tauschii*^25^ and *Ae. umbellulata*^26^ were downloaded, and redundant proteins were filtered with CD-HIT v4.8.1^27^ (parameters: -c 0.95). The resulted proteins were mapped against the PI 551049 assembly using GenomeThreader v1.7.1^28^. Finally, gene models from *Ae. tauschii* and *Ae. umbellulata* were transferred to the *Ae. comosa* genome using liftoff v1.6.3^29^ (parameters: -a 0.9 -s 0.9 -exclude_partial -polish). All outputs from the genome-guided and lifting methods were merged into a gff file using the Perl script “agat_sp_merge_annotation.pl” in the AGAT (https://github.com/NBISweden/AGAT). Proteins lacking start or stop codons, or shorter than 50 amino acids, were removed. The remaining protein sequences were compared against the UniProt database using BLASTP^30^ (parameters: -max_target_seqs 1 -outfmt 6 -evalue 1e-10). Redundant proteins from each gene were further removed using CD-HIT v.4.8.1^27^ (parameters: -c 1). The completeness of gene annotations was evaluated using BUSCO v1.7.1^31^. Finally, functional annotation of the protein-coding genes was performed using e eggNOG-mapper v2.1.12^32^, which assigns the functions to genes based on comparisons to the eggNOG v5.0.2 database.

### Synteny analysis

To explore syntenic blocks between *Ae. comosa* and other monocot/*Aegilops* species^26,33-35^, we identified collinear blocks based on protein sequences using JCVI v1.1.11^36^. Microcolinearity was also analyzed using JCVI v1.1.11.

### Investigation of *Ae. comosa* genome evolution

We performed phylogenetic analyses to determine the evolutionary relationships between *Ae. comosa* and 13 other diploid species. Orthologous genes among the 14 species were identified using OrthoFinder v2.5.4^37^. A total of 2,342 single-copy orthologous genes were identified and aligned using MUSCLE v5.1^38^, trimmed with trimAI v1.4.rev22^39^, and then concatenated for further analysis. A maximum likelihood phylogenetic tree was constructed based on the concatenated alignment using IQ-TREE v2.3.4^40^, with *B. distachyon* serving as the outgroup. To estimate divergence times, calibration points were obtained from the TimeTree database (www.timetree.org), specifically the *B. distachyon*–*H. vulgare* divergence (29.30–35.50 Mya) and the *H. vulgare*–*S. cereale* divergence (6.80–18.30 Mya). The CodeML and MCMCTREE programs withinthe PAML^41^ package were used to estimate divergence times among the 14 diploid species.

### Data Records

The PacBio HiFi reads, RNA-Seq, and Iso-seq data are available at the National Genomics Data Center, Beijing Institute of Genomics, Chinese Academy of Sciences, under BioProject accession number CRA016754 (https://ngdc.cncb.ac.cn/gsa/s/589tE69p). The genome assembly and annotation for PI 551049 were deposited in Figshare (https://figshare.com/s/55fe7056c3e03a15e39a; doi: 10.6084/m9.figshare.26928031).

### Technical Validation

We performed three key assessments of the assembled genome: (i) genome completeness using BUSCO^31^ v5.6.1 with the poales_odb10 lineage dataset; (ii) quality of the LTR assembly using LTR_FINDER_parallel^22^ v1.2 with the “-harvest_out” parameter and LTR_retriever^42^ v2.9.4 with the “-inharvest” option; and (iii) quality scoring using Merqury^43^ v1.3, based on NGS reads with a “k=21” parameter. The genome completeness, LTR assembly index (LAI), and consensus quality (QA) score of the *Ae. comosa* genome were 97.30%, 17.67, and 43.23, respectively, confirming the assembly as reference quality. Furthermore, the annotated proteins, evaluated with BUSCO using the poales_odb10 dataset, achieved a score of 97.60 %, highlighting the high quality of the annotation.

## Code availability

All software and pipelines were executed according to the manual and protocol of published tools. No custom code was generated for these analyses.

## Acknowledgements

This work was supported by the National Natural Science Foundation of China (32472159), the Key R&D Program of Shandong Province (ZR202211070163 and 2023LZGC022), the National Key Research and Development Program of China (2022YFD1201300), and the Young Taishan Scholars Program of Shandong Province.

## Author contributions

SC and LW conceived and supervised the study. HL generated the genome assembly and carried out the genome annotation. HL and SR analyzed the data and wrote the first version of the manuscript. RS collected the samples. RS, LQ, XH, JZ, KL, LH, and WH contributed to phenotyping, DNA extraction, and plant maintenance. SC obtained the funding and generated the final version of the manuscript. All authors have read and approved the final manuscript.

## Additional information

Correspondence and requests for materials should be addressed to S.C.

